# Age-Dependent, Odorant-Specific Changes in Olfactory Sensitivity in an Alzheimer’s Disease Mouse Model

**DOI:** 10.1101/2025.05.02.651821

**Authors:** Yuta Adachi, Kazuki Katori, Satoru Ishiyama, Shota Morikawa, Takashi Saito, Takaomi C. Saido, Yuji Ikegaya, Haruki Takeuchi

## Abstract

**Background:** Olfactory impairment often precedes cognitive decline in Alzheimer’s disease (AD). Recent studies suggest odor specificity in olfactory deficits during early AD stages, making olfactory tests a promising tool for early diagnosis. However, the mechanisms underlying olfactory impairment remain unclear, complicating the identification of optimal odorants for diagnostic purposes.

**Objective:** In this study, we assessed olfactory sensitivity in a knock-in mouse model of Alzheimer’s disease (*App^NL-G-F^* mice) that recapitulates key features of human AD pathology.

**Methods:** To evaluate odor detection thresholds, we employed an olfactory assay that leverages innate behavior without requiring associative learning. Six odorants representing distinct functional groups were tested in wild-type (WT) and *App^NL-G-F^* mice at 2 and 4 months of age.

**Results:** *App^NL-G-F^* mice exhibited odorant-specific hyposmia at 4 months of age, coinciding with amyloid deposition in cortical and subcortical regions but preceding measurable cognitive deficits. Unexpectedly, at an earlier stage (2 months), these mice showed odorant-specific hyperosmia to ester odorants, which transitioned to hyposmia by 4 months, indicating dynamic, age-dependent alterations in olfactory sensitivity as AD pathology progresses.

**Conclusions:** Our findings demonstrate that odorant-specific olfactory testing could serve as a promising diagnostic tool for early-stage AD, providing insights into the mechanisms underlying olfactory dysfunction in neurodegenerative diseases.

## Introduction

Alzheimer’s disease (AD) is the leading cause of dementia, affecting over 50 million people worldwide and projected to exceed 150 million cases by 2050, thus representing a significant social and economic burden in aging societies ^1,2^. Neuropathologically, AD is characterized by abnormal aggregation and deposition of amyloid-β (Aβ) protein as plaques, accompanied by neurofibrillary tangles consisting of hyperphosphorylated tau protein ^3–5^. These pathological hallmarks lead to progressive and irreversible neuronal damage, ultimately disrupting higher-order cognitive functions, including memory, language, visuospatial abilities, and executive control ^6–10^. Despite extensive efforts to develop pharmaceutical treatments targeting abnormal aggregation and deposition of Aβ, most therapeutic agents have failed to yield notable cognitive improvements during clinical trials. One hypothesis explaining these failures suggests that significant cognitive impairment correlates with advanced and possibly irreversible neuronal damage resulting from extensive Aβ and tau pathology. Consequently, research interest has increasingly shifted toward therapeutic strategies and interventions implemented at earlier pathological stages. Indeed, studies in AD mouse models have demonstrated that administration of antibodies targeting Aβ aggregation at the onset of plaque formation effectively suppresses further deposition, even one-month post-intervention ^11,12^.

Effective early intervention necessitates diagnostic tools capable of identifying AD pathology prior to noticeable cognitive decline. Recent developments in blood-based biomarkers, such as phosphorylated tau species, have gained attention for their diagnostic potential in preclinical AD stages; however, these approaches may require invasive sampling or complex laboratory infrastructure, potentially limiting widespread clinical applicability ^13–15^. In contrast, mounting evidence indicates that mild cognitive impairment (MCI)—an intermediate stage between normal cognition and AD—is frequently accompanied by olfactory dysfunction ^16–21^. Although the precise mechanisms remain under investigation, it is hypothesized that early amyloid-β or tau pathology in the olfactory bulb and related regions may disrupt olfactory processing. Because olfaction can be assessed using non-invasive, cost-effective methods that require only minimal infrastructure, olfactory testing emerges as a particularly promising and complementary approach for the early detection and diagnosis of AD.

Animal models have been extensively utilized to investigate the pathological and behavioral manifestations of Alzheimer’s disease (AD). Among these, mouse models are the most widely employed not only for evaluating the efficacy of potential therapeutic interventions but also for elucidating the underlying mechanisms of AD pathology ^22–24^. However, transgenic mouse models of AD often overexpress amyloid precursor protein (APP), which may lead to artificial phenotypes not representative of human AD pathology. Saito et al. developed a knock-in AD mouse model carrying three mutations— the Swedish, Iberian, and Arctic—that are known to cause familial AD. The mice have been shown to exhibit Aβ deposition in the brain without overexpression of APP ^25^. While this AD mouse model (*App^NL-G-F^* mouse) is useful for studying how Aβ amyloidosis induces subsequent pathophysiological changes, its potential for investigating olfactory function remains to be fully elucidated.

In this study, we utilized an automated olfactory test that leverages innate olfactory behaviors to assess odor detection deficits in *App^NL-G-F^* mice. We observed odorant-specific hyposmia at 4 months of age, coinciding with the onset of cortical and subcortical amyloidosis but preceding measurable cognitive decline. Additionally, we found odorant-specific hyperosmia at an earlier stage (2 months), when amyloid plaques are not yet detectable. These observations not only demonstrate the usefulness of the knock-in AD mouse model for studying early olfactory deficits characteristic of human AD but also suggest that odorant-specific olfactory testing may serve as a novel early diagnostic marker for the disease.

## Materials and Methods

### Animals

All experiment procedures were performed with the approval of the animal experiment ethics committee of the University of Tokyo. *App^NL-G-F^*mice carrying Arctic, Swedish, and Iberian mutations were provided by the RIKEN BRC. C57BL/6J mice, purchased from SLC (Japan), were used as wild-type (control) mice. The wild-type mice were only male, whereas the *App^NL-G-F^* mice were of both sexes. Mice were housed under a 12-hour light/dark cycle and had ad libitum access to water and food.

### Odors

Odorants were purchased from Tokyo Chemical Industry Co. Each odorant was diluted in mineral oil (MO; Nacalai Tesque), and 200 µl was placed in an odor bottle every experimental day. Odorants used in this study included isoamyl acetate, 2-acethyl-5-metylfuran, 2-heptanone, citronellal, limonene, eugenol, and ethyl butyrate. These odorants were chosen based on previous research using various functional groups ^26,27^.

### Odor detection threshold test

The experiments were conducted using mice aged 9-11 weeks (designated as the 2-months group) or 17-19 weeks (designated as the 4-months group). Each experimental group included 6-15 animals. All animals were tested only once per day during their light phase.

Odor detection threshold tests were conducted in polyvinyl chloride (PVC) boxes (200 × 200 × 200 mm), each featuring an odor delivery port on one wall. A custom-made olfactometer regulated the timing and delivery of odor stimuli using solenoid valves controlled by an Arduino and custom-written scripts. The air from the air pump was divided into two paths, one to a 3-way solenoid valve and the other to a 2-way solenoid valve. The 3-way solenoid valve managed the primary airflow and control stimulus (mineral oil) delivery. During baseline periods and inter-trial intervals, the normally open (NO) port of this valve delivered a blank air stream at 300 mL/min into the test box; the normally closed (NC) port remained closed during these periods. The NC port was connected to a bottle containing mineral oil. For control trials (mineral oil presentation), the 3-way valve switched actuation, closing the NO port and simultaneously opening the NC port to deliver air passed through the mineral oil bottle at 400 mL/min. A separate 2-way solenoid valve, connected to a bottle containing the odorant, controlled odorant delivery. During odor presentation trials, the 3-way valve remained in its normal state (NO port delivering 300 mL/min blank air), and the 2-way valve opened to add a 100 mL/min air stream passed through the odorant bottle. This odorized stream merged with the blank air from the NO port, resulting in a total airflow of 400 mL/min. To minimize residual odors, air was continuously evacuated from the box via an exhaust tube located on the wall opposite the odor delivery port, connected to a vacuum pump.

Each trial consisted of a 2-minute baseline period with blank air delivery, followed by a 1-minute stimulus presentation period. Animals first underwent seven trials where mineral oil was presented during the stimulus period to establish baseline nose-poking duration. In the 8th trial, the odorant of interest was presented. Odor investigation behavior was quantified by recording the duration of nose pokes into the odor port, detected by an infrared sensor. Data acquisition was performed using the same software that controlled the olfactometer.

To quantify the odor investigation behavior relative to a baseline, we defined a normalized port investigation (NPI) for the i^th^ trial as follows:

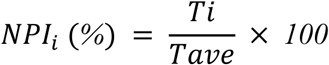

where *T*i represents the duration of the mouse’s nose-poking into the odor port during the i^th^ trial, and *T*ave is the average baseline nose-poking duration. *T*ave was calculated using the data from the first five baseline trials:

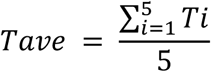

To quantify the change in exploratory behavior specifically elicited by the odor presentation, we defined ΔNPI as the difference between the NPI during the odor trial (trial 8) and the NPI during the final baseline trial (trial 7):

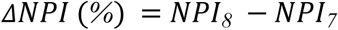

The odorant was defined as detectable if the ΔNPI value was significantly greater than 0.

### Methimazole administration

Methimazole (1-methyl-2-mercaptoimidazol) has been reported to induce apoptotic cell death of olfactory neurons in mice, resulting in anosmia at least two to six days after administration ^28,29^. Therefore, to ablate OSNs, mice were injected intraperitoneally with methimazole (75 mg/kg; FUJIFILM Wako Pure Chemical Corporation) dissolved in 0.9% saline 4 days before the behavioral experiment. Control mice were injected intraperitoneally with 0.9% saline.

### Y-maze test

The Y-maze apparatus (O’Hara & Co) was made of gray PVC and had three compartments (3 cm bottom, 12 cm top, 40 cm long, and 12 cm high) radiating from a central platform (3 x 3 x 3 cm triangle). The light intensity at the center of the maze was maintained at 20 lx. The Y-maze test was conducted during the light. Briefly, each mouse was initially placed in the center of the maze, facing one of the arms, and allowed to freely explore the maze for 8 minutes. The activity of the mice was recorded using a web camera (Logicool) positioned above the maze. The position of the mouse was determined based on the head and tail-base, which were estimated using DeepLabCut ^30^. An arm entry was recorded when both the head and tail-base of the mouse crossed the border defining the arm entrance. Spontaneous alternation behavior was quantified using the alternation score, calculated as follows:

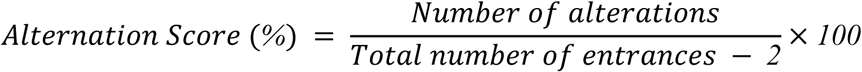

An ‘alteration’ was defined as a sequence of three consecutive entries into distinct arms (e.g., visiting arm A, then B, then C constitutes one alteration). The total number of possible alternations is the total number of arm entries minus two.

### Quantification and statistical analysis

All the statistics are conducted in MATLAB. Data were shown as means ± SEMs in figures and text. Significance was defined as: * indicates p<0.05, ** indicates p<0.01, *** indicates p<0.001, n.s. indicates not significant.

## Result

Conventional olfactory tests, such as the Go/No-Go task or forced choice task, have been widely used to assess odor detection thresholds ^31^. However, these behavioral assays rely on associative learning, making it difficult to determine whether observed deficits in AD model mice are due to olfactory dysfunction or learning abnormalities. To overcome this limitation, we employed an olfactory behavioral test that leverages innate olfactory behaviors, wherein mice exhibit curiosity-driven exploration of novel odors ^32^. This method avoids the confounding effects of cognitive impairments, allowing for a more precise assessment of odor detection thresholds. In this test, a mouse was placed in a behavioral chamber equipped with a single odor delivery port, and the duration of nose pokes during odor presentations was measured (Fig. 1A). To establish baseline exploration behavior, mineral oil was delivered for 1 minute, interspersed with 2-minute intervals, and the nose-poke duration was recorded. After seven trials with mineral oil, the test odorant was presented for 1 minute. The duration of nose pokes during odor exposure was normalized to the baseline established with mineral oil as Normalized Port Investigation (NPI). If the presented odor concentration was sufficient for detection, mice showed an increase in nose-poke durations compared to the last (7th) mineral oil exposure (Fig. 1B). Anosmic mice induced by methimazole treatment, which ablates olfactory sensory neurons ^33^, did not exhibit increased nose-poking behavior, confirming that the test is olfactory-dependent (Fig. 1C).

**Fig 1.**
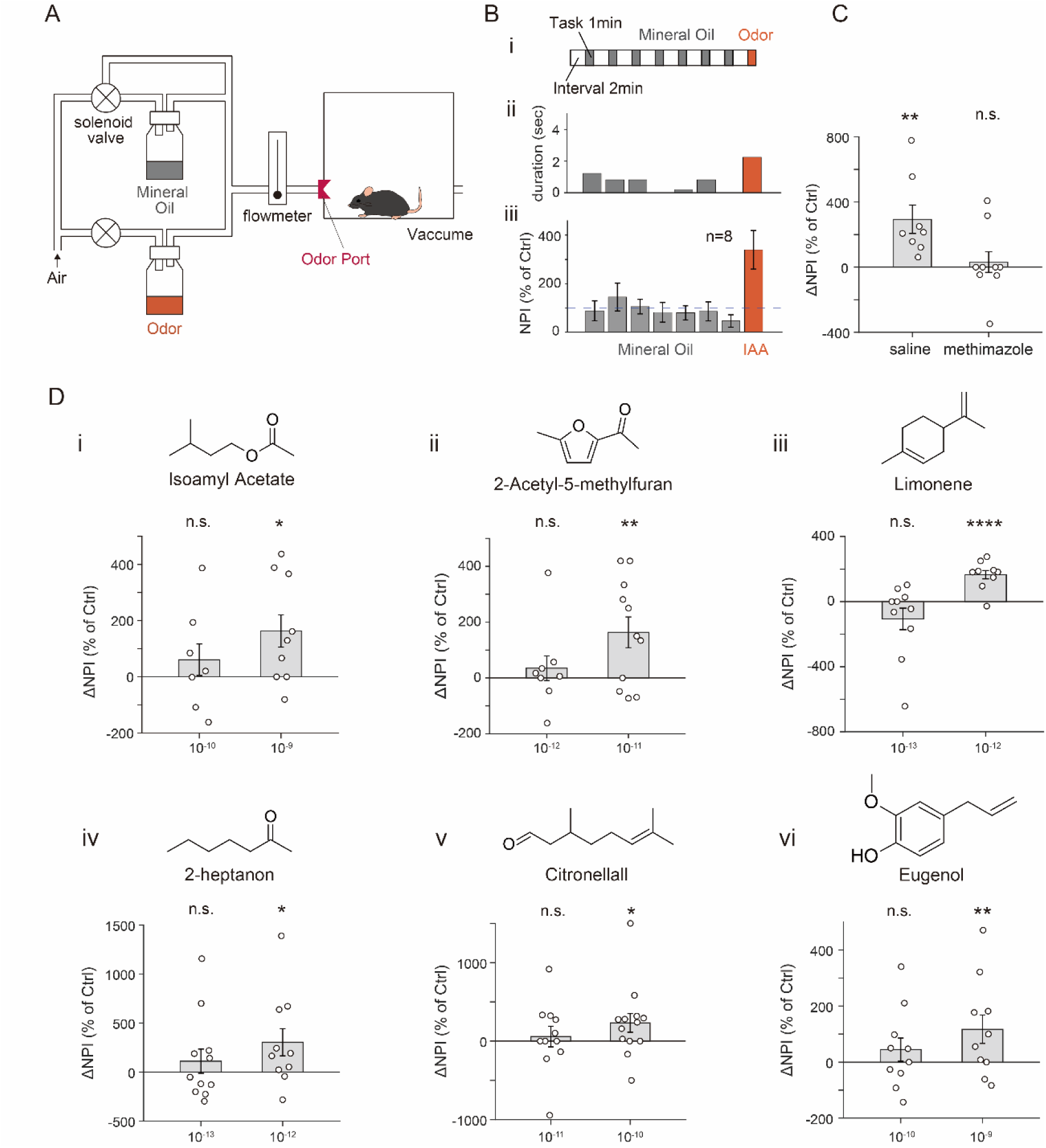
Olfactory behavioral test for determining odor threshold concentration in WT mice. A) A schematic diagram of the apparatus used to measure the odor detection threshold of mice. Solenoid valves control the supply of air containing either mineral oil (MO, gray) or odorant (orange) through a port in the cage. B) i. Schematic showing odor delivery sequence in the odor detection test. Mineral Oil (gray) or odor (orange) was presented for 1 minute with a 2-minute interval. ii. A representative bar plot showing the nose-poke duration of a single mouse. The odor used was Isoamyl Acetate (10^-6^ (v/v)). iii. Bar plot of average NPI. (*n* = 8 mice) C) Bar plot of average ΔNPI of control (saline) mice and OSN-ablated (methimazole) mice (*n* = 8 for control mice, *n* = 9 for OSN-ablated mice). D) Bar plots of ΔNPI for six odorants at the threshold concentration (right) and ten-fold below (left). The structural formula of each odorant is shown at the top. The following odorants were used. Isoamyl Acetate (10⁻⁹ (v/v), *n* = 7; 10⁻⁸ (v/v), *n* = 9), 2-Acetyl-5-methylfuran (10⁻¹² (v/v), *n* = 8; 10⁻¹¹ (v/v), *n* = 11), Limonene (10⁻¹³ (v/v), *n* = 10; 10⁻¹² (v/v), *n* = 9), 2-Heptanone (10⁻¹³ (v/v), *n* = 11; 10⁻¹² (v/v), *n* = 10), Citronellal (10⁻¹¹ (v/v), *n* = 13; 10⁻¹⁰ (v/v), *n* = 11), and Eugenol (10⁻¹⁰ (v/v), *n* = 10; 10⁻⁹ (v/v), *n* = 10). Data are presented as scatter plot for individual values, with bar plots representing the mean ± SEM. Pairwise t-test p values are shown. *P<0.05, **P<0.01, ***P<0.0001, n.s., not significant.

Using this system, we determined the detection threshold concentrations of six neutral odorants—isoamyl acetate (Esters), 2-acetyl-5-methylfuran (Furans), 2-heptanone (Ketones), citronellal (Aldehydes), limonene (Terpenes), and eugenol (Vanillins)—each representing a distinct functional group, in WT mice. To determine the detection threshold, we identified the minimum concentration at which mice could reliably detect the odor compared to mineral oil ^32^. For this, odorants were first presented at low concentrations, and if mice did not respond, the same odor was presented at a ten-fold higher concentration on the following day. This procedure continued until mice showed a significant increase in nose-poking behavior (Fig. 1D). We found that the olfactory detection thresholds for WT mice to Isoamyl Acetate (IAA), 2-Acetyl-5-methylfuran, Limonene, 2-heptanone, Citronellal, and Eugenol were 10^-8^, 10^-11^, 10^-12^, 10^-12^, 10^-10^, and 10^-9^ (v/v), respectively.

Next, we conducted similar experiments using Alzheimer’s disease (AD) model mice. Specifically, we used *App^NL-G-F^* mice, which carry three familial Alzheimer’s disease-associated mutations: the Swedish mutation, Beyreuther/Iberian mutation, and the Arctic mutation ^25^. These mice are known to develop cortical amyloid deposition by 2 months of age, with amyloidosis in the subcortical regions typically beginning around 4 months, while cognitive behavioral abnormalities are not observed until 6 months ^25^. To validate the odor detection threshold in mice, we conducted an olfactory threshold test using a different cohort of WT mice from those used in Figure 1. The test was performed at the threshold concentration and a concentration ten times lower. The results confirmed that the mice responded exclusively to the odorant at the threshold concentration. Next, we examined whether 4-month-old *App^NL-G-F^* mice, at an age when cortical amyloidosis is present without cognitive deficit, responded to the odorant at the threshold concentration (Fig. 2A-F). Among the six tested odorants, we found that *App^NL-G-F^* mice did not respond to four odorants—IAA, 2-Acetyl-5-methylfuran, Limonene, and Citronellal—at the threshold concentrations established for WT mice (IAA: 10^-8^, 2-Acetyl-5-methylfuran: 10^-11^, Limonene: 10^-12^, Citronellal: 10^-10^ (v/v)). Consistent with a previous finding ^25^, no significant cognitive impairments were observed in these 4-month-old *App^NL-G-F^* mice in the Y-maze test (Fig. 2G). These results suggest that olfactory impairment precedes cognitive decline in *App^NL-G-F^*mice, mirroring the early progression of human Alzheimer’s disease (AD).

**Fig 2.**
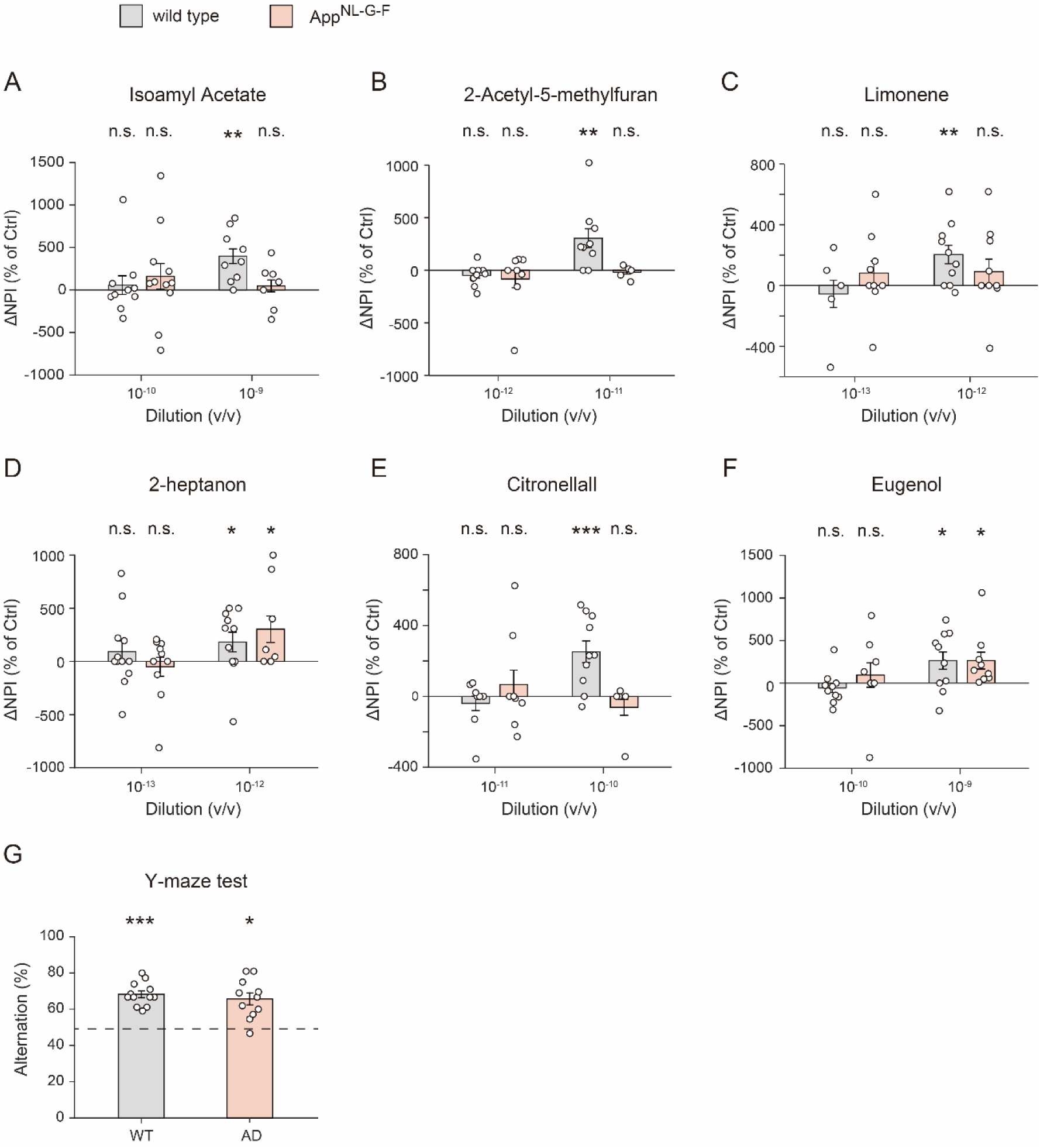
Hyposmia to specific odorants in 4-month-old *App^NL-G-F^* mice. A-F) Average ΔNPI for wild-type mice (WT, gray) and 4-month-old *App^NL-G-F^* mice (orange) across six odorants at the threshold concentration (right) and ten-fold lower concentrations (left). The following odorants were used. Isoamyl Acetate (10⁻⁹ (v/v), WT *n* = 10, *App^NL-G-F^ n* = 11; 10⁻⁸ (v/v), WT *n* = 9, *App^NL-G-F^ n* = 7), 2-Acetyl-5-methylfuran (10⁻¹² (v/v), WT *n* = 9, *App^NL-G-F^ n* = 8; 10⁻¹¹ (v/v), WT *n* = 9, *App^NL-G-F^ n* = 6), Limonene (10⁻¹³ (v/v), WT *n* = 5, *App^NL-G-F^ n* = 9; 10⁻¹² (v/v), WT *n* = 10, *App^NL-G-F^ n* = 9), 2-Heptanone (10⁻¹³ (v/v), WT *n* = 12, *App^NL-G-F^ n* = 10; 10⁻¹² (v/v), WT *n* = 11, *App^NL-G-F^ n* = 7), Citronellal (10⁻¹¹ (v/v), WT *n* = 8, *App^NL-G-F^ n* = 8; 10⁻¹⁰ (v/v), WT *n* = 10, *App^NL-G-F^ n* = 5), and Eugenol (10⁻¹⁰ (v/v), WT *n* = 10, *App^NL-G-F^ n* = 7; 10⁻⁹ (v/v), WT *n* = 10, *App^NL-G-F^ n* = 9). G) Percentage of spontaneous alternation in the Y-maze for wild-type (WT) and 4-month-old *App^NL-G-F^* mice (*n* = 12 for WT mice, *n* = 11 for *App^NL-G-F^* mice). Data are presented as scatter plot for individual values, with bar plots representing the mean ± SEM. Pairwise t-test p values are shown. *P<0.05, **P<0.01, ***P<0.005, n.s., not significant. Pairwise t-test p values are shown. *P<0.05, **P<0.01, ***P<0.005, n.s., not significant.

We then tested the 2-month-old *App^NL-G-F^* mice, at an age when cortical amyloid deposition begins. *App^NL-G-F^* mice responded to all four odorants tested at the threshold concentrations (Fig. 3A-D). However, unexpectedly, they demonstrated heightened sensitivity to IAA, responding at a concentration ten times lower than the threshold established for WT mice (Fig 3A). This finding suggests that hyperosmia to specific odorants may emerge during the early stages of AD. Given that IAA belongs to the ester group, we further tested another ester compound, ethyl butyrate. Interestingly, as observed in the experiment with IAA, 4-month-old *App^NL-G-F^* mice failed to respond to ethyl butyrate at the threshold concentration determined by WT mice, while 2-month-old *App^NL-G-F^*mice responded at a concentration ten times lower than the threshold. These findings demonstrate that the olfactory sensitivity of AD model mice to ester compounds dynamically changes with the progression of AD pathology. Specifically, 2-month-old *App^NL-G-F^* mice exhibit heightened sensitivity, whereas 4-month-old *App^NL-G-F^* mice demonstrate reduced sensitivity. Importantly, these changes in olfactory thresholds occur before the onset of cognitive impairments, suggesting that olfactory function could serve as a diagnostic marker for early AD.

**Fig 3.**
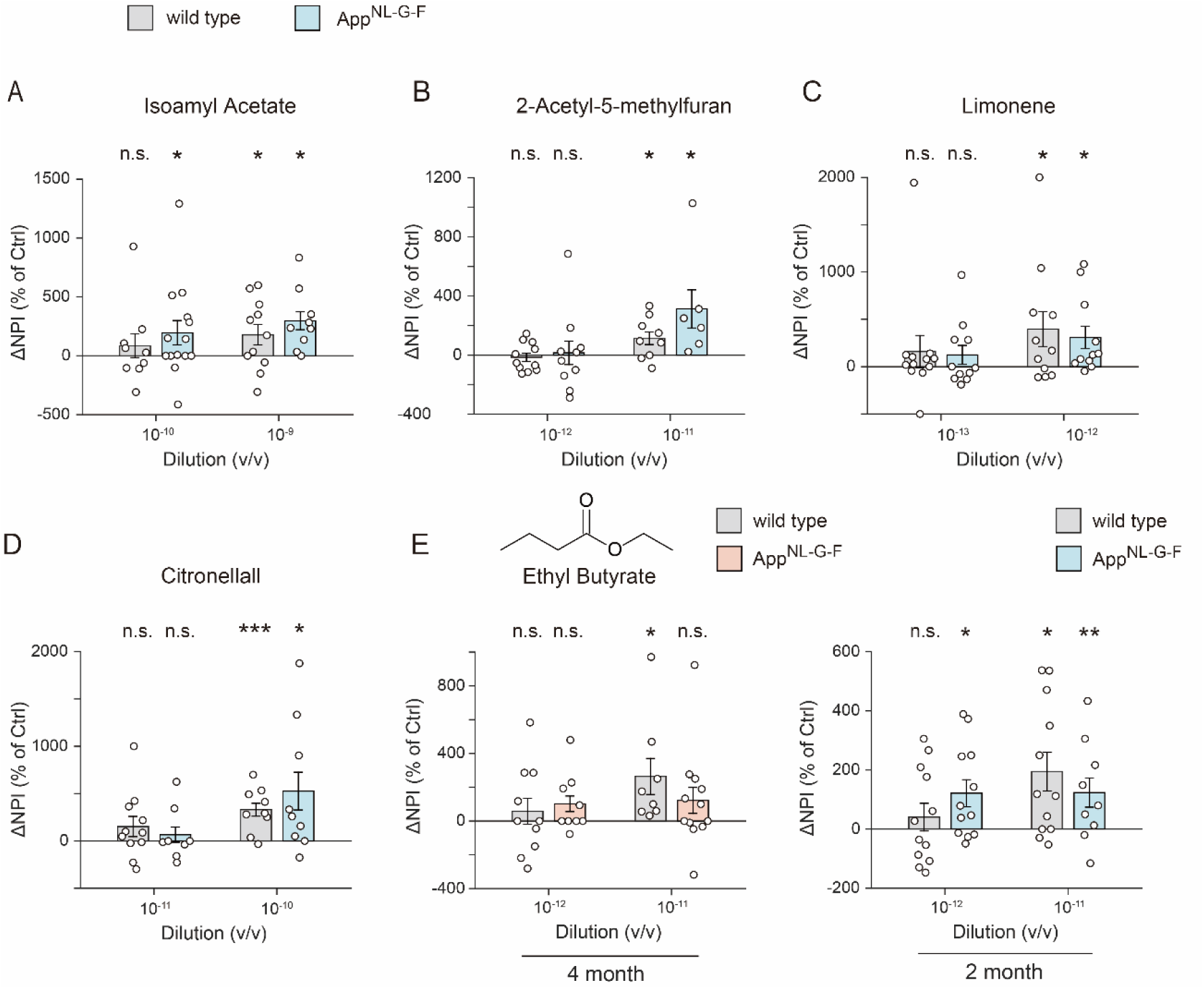
Increased olfactory sensitivity to esters in 2-month-old *App^NL-G-F^* mice. A-D) Average ΔNPI for wild-type mice (WT, gray) and 2-month-old *App^NL-G-F^*mice (blue) across four odorants at the threshold concentration (right) and ten-fold lower concentrations (left). The following odorants were used. Isoamyl Acetate (10⁻⁹ (v/v), WT *n* = 9, *App^NL-G-F^ n* = 14; 10⁻⁸ (v/v), WT *n* = 11, *App^NL-G-F^ n* = 9), 2-Acetyl-5-methylfuran (10⁻¹² (v/v), WT *n* = 11, *App^NL-G-F^ n* = 10; 10⁻¹¹ (v/v), WT *n* = 9, *App^NL-G-F^ n* = 6), Limonene (10⁻¹³ (v/v), WT *n* = 12, *App^NL-G-F^ n* = 11; 10⁻¹² (v/v), WT *n* = 11, *App^NL-G-F^ n* = 11), and Citronellal (10⁻¹¹ (v/v), WT *n* = 11, *App^NL-G-F^ n* = 8; 10⁻¹⁰ (v/v), WT *n* = 9, *App^NL-G-F^ n* = 9). E) Bar plots of wild-type mice (gray) and *App^NL-G-F^*mice (orange, or blue) for Ethyl Butyrate. The left plot shows data from 4-month-old mice (10^-12^ (v/v), WT *n* = 10, *App^NL-G-F^ n* = 9, 10^-11^ (v/v), WT *n* = 8, *App^NL-G-F^ n* = 12) and the right plot from 2-month-old mice (10^-12^ (v/v), WT *n* = 12, *App^NL-G-F^ n* = 12, 10^-11^ (v/v), WT *n* = 12, *App^NL-G-F^ n* = 9). Data are presented as scatter plot for individual values, with bar plots representing the mean ± SEM. Pairwise t-test p values are shown. *P<0.05, **P<0.01, ***P<0.005, n.s., not significant.

## Discussion

Olfactory impairment is widely recognized as an early symptom of AD, indicating the potential use of olfaction as an early diagnostic marker. Because olfactory function generally declines with age ^34^, it is crucial to develop measurement methods— encompassing both the selection of odorant combinations and the assessment of olfactory function—when applying olfactory testing for AD diagnosis.

Animal models have proven extremely useful for investigating Aβ pathology ^35^. However, previous studies on olfactory function using AD model mice have often been limited by the small number of odorants tested or the lack of systematic evaluation of odorant concentrations. In many cases, studies have focused merely on confirming whether olfactory function was present or absent, rather than conducting a thorough examination of olfactory thresholds^16,36,37^. Additionally, AD models that rely on overexpression of amyloid precursor protein (APP) may introduce artifacts associated with non-physiological APP levels ^25,35^. In the present study, we employed a next-generation AD model mouse that more closely mirrors human Aβ pathology. Using six odorants representing different functional groups, each tested at multiple concentrations, we conducted olfactory threshold assays. Our results demonstrated that certain odorants exhibited hyposmia before obvious cognitive deficits. Moreover, we found that at an earlier stage—when no amyloid plaques were yet detectable in the brain—the mice displayed hyperosmia specifically toward ester compounds. Numerous studies have investigated abnormal network excitability in AD ^38–41^, and moderate levels of Aβ are thought to induce neuronal hyperexcitability ^42^. Consistent with this, Wesson et al. (2011) observed hyperactive odor-evoked responses in the PC and increased functional connectivity between the OB and PC in Tg2576 mice aged 6–7 months, a stage at which Aβ deposition was modest ^43^. Together, these findings suggest that moderate Aβ accumulation may lead to odor hypersensitivity, which subsequently transitions to hyposmia as pathology advances.

One noteworthy aspect of this study is the observation of odor-specific deficits in olfactory function. Notably, such odor-specific impairments have also been reported in human clinical studies ^17,44,45^. Because Aβ accumulation has been reported throughout the olfactory system—from peripheral to central structures—it remains unclear which region is primarily responsible for the odor-specific impairments observed in AD ^46–48^. Odor information detected by peripheral sensory neurons is relayed to the olfactory bulb (OB), where it is transformed into a topographic odor map; in this map, chemical features of odorants are spatially organized ^49–52^. In contrast, higher olfactory areas such as the piriform and entorhinal cortices largely discard this spatial organization, representing odor information instead through distributed and non-topographic patterns of neuronal activity ^53–55^. If central olfactory regions were significantly impaired, one would expect general olfactory deficit, rather than selective impairments for specific odorants. Therefore, the odor-specific deficits observed in this study are more likely to reflect abnormalities in the OB or even more peripheral structures, such as the olfactory epithelium (OE). Indeed, several studies have reported pathological alterations in the OE of AD model mice ^56^. For example, in 5×FAD mice, region-specific Aβ accumulation was observed in the OE, suggesting that certain peripheral zones may be more vulnerable to early pathology. While tau pathology in higher-order brain regions and disruption of cholinergic input are also considered potential contributors to olfactory dysfunction in AD ^57,58^, these mechanisms are more likely to result in generalized rather than odor-specific deficits. Further investigation is needed to determine whether the olfactory impairments observed in *App^NL-G-F^* mice arise from peripheral or central changes, and whether similar mechanisms underlie the olfactory deficits seen in human AD.

## Acknowledgements

We thank T. Kimura for her help in preparing the manuscript.

## Author Contributions

Yuta Adachi (Conceptualization; Data Curation; Formal analysis; Investigation; Methodology; Visualization; Writing – original draft, Writing – review & editing), Kazuki Katori (Conceptualization; Data Curation; Formal analysis; Funding acquisition; Investigation; Methodology; Supervision; Visualization; Writing – original draft, Writing – review & editing), Satoru Ishiyama (Investigation; Resources; Visualization, Writing – original draft), Shota Morikawa (Resources; Supervision), Takashi Saito (Resources), Takaomi C. Saido (Resources), Yuji Ikegaya (Resources; Supervision), Haruki Takeuchi (Conceptualization; Funding acquisition; Project administration; Supervision; Writing – original draft, Writing – review & editing)

## Statements and declarations

Declaration of conflicting interests

The authors declare no potential conflicts of interest with respect to the research, authorship, and publication of this article.

## Data availability

The data supporting the findings of this study are available on request from the corresponding author.

